# Infection of neotropical non-human primates with bat deltaviruses

**DOI:** 10.64898/2026.09.02.747548

**Authors:** Felix Lehmann, Andres Moreira-Soto, Maria Angélica Mares-Guia, Alejandro Alfaro-Alarcón, Camila Vieira Molina, Ianei Oliveira Carneiro, Breno Frederico de Carvalho Dominguez Souza, Aroldo José Borges Carneiro, Gabriela Hernández-Mora, Dieter Glebe, Angélica Cristine de Almeida Campos, Carlos Roberto Franke, Edison Luiz Durigon, Ana Maria Bispo de Filippis, Jan Felix Drexler

## Abstract

**Background & Aims:** The origin of hepatitis D virus (HDV) and related deltaviruses remains elusive. Contrarily, hepatitis B virus, HDV’s helper virus, and related hepadnaviruses display long-term association with primates. Current data suggest cross-order host shifts as a common mechanism in deltavirus ecology, but corroborative evidence is lacking. Here, we aimed to elucidate the genealogy of primate deltaviruses.

**Methods:** We screened 961 non-human primate (NHP) liver specimens obtained in Brazil between 2017-2023 for deltaviruses and hepadnaviruses. Complete deltaviral genomes were obtained via overlapping nested RT-PCR, cloned, expressed *in vitro* and analyzed via immunoblot and immunofluorescence analysis. Anti-deltaviral antibodies in NHP and vampire bats were detected via immunofluorescence analysis.

**Results:** We detected deltaviruses in two NHP. Complete deltaviral genomes exhibited common features including high self-complementarity, genomic and antigenomic ribozymes, and a delta antigen open reading frame. NHP deltaviruses were phylogenetically related to viruses found in common vampire bats. The NHP deltaviruses replicated *in vitro* without the expression of a large delta antigen. We did not detect anti-deltaviral antibodies in NHP sera (0/249), in contrast to sera from common vampire bats (7/112; 6.2%, 95% CI: 1.8-10.7), indicating viral circulation in bats. Ancestral state reconstruction suggested a bat origin of NHP deltaviruses. Targeted screening excluded a coinfecting hepadnavirus in the deltavirus-positive animals but led to the discovery of a hepadnavirus, corroborating a non-recent introduction of hepadnaviruses into the primate stem-lineage.

**Conclusions:** Our data are consistent with deltaviral cross-order host shifts and suggestive of the susceptibility of primates to reservoir-bound deltaviruses, lending credibility to a zoonotic origin of HDV.

## INTRODUCTION

The evolutionary history of the human hepatitis D virus (HDV) and related non-human deltaviruses (family *Kolmioviridae*) remains puzzling, mainly due to the lack of clear virus-host co-evolution, potentially attributable to frequent transmissions between divergent hosts. The occurrence of such host shifts is likely facilitated by the ability of deltaviruses to autonomously replicate in a diverse array of cell types from different animals [1].

All known deltaviruses only encode a single protein, the delta antigen (DAg, for HDV: S-DAg), required for viral replication [2]. At present, HDV is the only deltavirus known to express a second, C-terminally extended isoform of DAg (large DAg, L-DAg) generated during late-stage replication by site-specific mutation of the DAg amber stop codon mediated by the cellular adenosine deaminase acting on RNA 1 (ADAR1) protein. The expression of L-DAg marks the switch from genome replication to viral egress, as the extension of L-DAg binds to the surface proteins of hepatitis B virus (HBV), HDV’s exclusive helper virus *in vivo*, on which HDV depends for envelopment and secretion of infectious viral particles [3]. Contrarily, envelopment of non-human deltaviruses may occur through opportunistic packaging into budding particles of different evolutionarily independent viruses, a mechanism recently likened to the Trojan horse [4], explaining the seemingly arbitrary host and tissue tropisms of non-human deltaviruses observed.

Due to HBV’s strong hepatotropism, HDV is also limited to the liver, where it causes the most severe form of chronic viral hepatitis [5]. Data on pathogenesis of non-human deltaviruses are currently limited to a single study of the Boa constrictor deltavirus (bconDV), where bconDV-positive tissues (i.e., brain, liver, lung, spleen and kidney) did not show pathohistological changes attributable to deltavirus infection [6]. At present, the absence of *bona fide* deltavirus/helper virus pairings impedes studies on their pathogenesis, which may differ among infected host species and tissues.

Lastly, a New World origin of mammalian deltaviruses is the currently favored hypothesis, as most non-human deltaviruses have been detected in the New World despite substantial bias towards the Old World in terms of diversity of species analyzed and sequence datasets available [7]. Conclusive evidence on when and where humans potentially acquired the virus ancestral to HDV, and whether its unique dependence on a hepadnavirus (like HBV) through means of the L-DAg arose prior to or after this introduction, remains elusive in the absence of known animal reservoirs. Therefore, targeted screening of mammals in South America, a global hotspot of faunal diversity [8], is vital to understand the complex genealogy of mammalian deltaviruses. Here, we describe the discovery and characterization of two novel deltaviruses discovered in Neotropical primates and discuss their implications for deltavirus ecology.

## MATERIAL AND METHODS

### Sampling

Liver specimens were obtained between 2017-2023 from deceased wild non-human primates (NHP) in Brazil conducted by the Oswaldo Cruz Foundation (Fiocruz) in Rio de Janeiro (resolution 2.998.362 IOC/Fiocruz; CITES permit no.: 23BR046601/DF). NHP sera samples used for serology were obtained in Brazil by the Institut Pasteur de São Paulo (Bioethics Committee of the ICB/USP Protocol No. 3971161219; CITES permit no.: 22BR043788/DF) or previously described [9]. Sera and organ specimens from common vampire bats (*Desmodus rotundus*) from Brazil and Costa Rica used for deltavirus screening and/or serological testing were previously described [10].

### Screening and genome amplification

Nucleic acids were isolated using the “MagNA Pure 96 DNA and Viral NA Small Volume Kit” on the MagNA Pure 96 instrument (Roche). Samples were subjected to a hepadnavirus-[11] or deltavirus-specific nested PCR using degenerate primers located in the S and DAg open reading frame (ORF), respectively (**Supplementary CTAT Table**). Viral genomes were completed using sequence-specific primers (**Supplementary CTAT Table**) and reaction products were Sanger sequenced (Microsynth). Realtime RT-PCR to determine the viral loads of organ specimens of K083_DR135 included a standard based on *in vitro* transcribed RNA of a synthetic gene fragment (Integrated DNA technologies) covering the amplicon region.

### Bioinformatics

Full-length nucleotide (hepadnaviruses) or DAg amino acid sequences (deltaviruses) were aligned using MAFFT implemented in Geneious Prime. Bayesian phylogenies were obtained using MrBayes [12] with 2 million generations, sampling every 100^th^ generation and a burn-in of 25%. An HKY+G substitution model (as described in [11]) and a WAG substitution model were employed for nucleotide and amino acid alignments, respectively. Ancestral state reconstructions (ASR) were run for 10 million generations, sampling every 1,000^th^ generation and a burn-in of 25%. Nucleotide and amino acid distances were based on MAFFT alignments and calculated in MEGA X with pairwise deletion option [13]. RNA folding structures and nucleotide complementarity plots were generated using the mfold webserver [14]. ORFs were predicted in Geneious Prime using the “Find ORFs” feature setting a minimum size of 200 nucleotides and ATG start codon. Transmembrane domains and signal peptides were predicted using the DeepTMHMM 1.0 and SignalP 6.0 webservers, respectively [15,16].

### Plasmids

The complete ORFs of the DAg of the novel deltaviruses were synthesized and cloned into the multiple cloning site of pcDNA3.1(+) (Biocat). Cloning of deltaviral genomic head-to-tail dimers into pcDNA3.1(+) was performed as described [17] using sequence-specific primers (**Supplementary CTAT Table**). Dimers containing an R13A mutation within the DAg ORF (abolishing antigenome to genome conversion in HDV [18], **Fig. S1A**) in both copies of the genome were generated via site-directed mutagenesis and verified through Nanopore sequencing (Microsynth). The R13A mutation was also introduced into a plasmid carrying a human HDV-1 genomic dimer (GenBank accession number: KY495779).

### Cell culture

The human hepatoma cell line HuH7 was cultured in complete DMEM (high-glucose DMEM with 10% fetal bovine serum (FBS), 100 U/mL penicillin, and 100 μg/mL streptomycin) with 5% CO_2_ at 37°C. Contamination with mycoplasma was excluded through quantitative PCR (Eurofins Genomics).

### Cell division-mediated spread

HuH7 cells were transfected with wildtype and R13A deltavirus dimer plasmids using FugeneHD (Promega, #E2311) following the manufacturer’s instructions, with 250 ng plasmid DNA per cm^2^ of cells at 80% confluence and a DNA:reagent ratio of 4:1 in OptiPRO SFM (Gibco, #12309050). The following day, cells were detached using trypsin-EDTA 0.05% (Gibco, #25300054), split 1:100 and cultivated in complete DMEM until 10 days post transfection (dpt) with media changes every two days. At 10 dpt, cells were washed with PBS and fixed/permeabilized using ROTI-Histofix (Carl Roth, #P087.4) with 0.2% Triton-X 100 (15 min, RT). Cells were blocked using 10% FBS in PBS for 1h at RT and then probed with primary antibody (anti-Proechimys semispinosus deltavirus (psemDV) rabbit serum (SY0041, generated in [17], **Supplementary CTAT Table**) in PBS; 1:400 dil.; 1 h, 37°C). After washing three times with PBS, cells were incubated with secondary antibody (Cy3-coupled goat anti-rabbit IgG in 50% glycerol, 1:200 dil.) and 1 µg/ml DAPI (1h, RT). After three washes, cells were overlayed with PBS for storage. Image acquisition was performed at 100 × or 200 × total magnification on a DMi8 inverse microscope and a DFC9000 GTC camera (both Leica) using the Leica Application Suite X.

### Immunoblot

HuH7 cells were transfected as described above. Starting from 1 dpt, medium (complete DMEM with 1% DMSO) was exchanged every two days until 10 dpt. At 4, 6, 8, and 10 dpt, one well was harvested as follows: Cells were washed with PBS and then 100 µl M-PER (Thermo Scientific, #78501) per one well of a 24-well plate were added (5 min, RT). The suspensions were transferred to reaction tubes and centrifuged (5 min, 16,000 × g). The supernatant was transferred to new reaction tubes and stored at −20°C until use. Cell lysates were mixed with 4 × loading buffer (0.2 M tris-HCl, 0.4 M DTT, 277 mM SDS, 6 mM bromophenol blue, 4.3 M glycerol in water) and subjected to discontinuous SDS-PAGE (14% separating gel). Separated protein was semi-dry blotted onto methanol-activated 0.45 µm PVDF membranes (Thermo Scientific, #88518) using Bjerrum Schafer-Nielsen buffer (40 mM tris, 30 mM glycine in water with 20% v/v methanol freshly-added) at 25 V for 45 min. Membranes were blocked (1 h, RT) with 5% w/v non-fat milk powder in TBS-T (0.1% Tween-20). After blocking, membranes were cut below the 40 kDa marker band for individual antibody incubation. Membranes were probed (overnight, 4°C) with primary antibody in 1% w/v non-fat milk powder in TBS-T with 0.02% sodium azide (either 1:2,000 dil. anti-psemDV rabbit serum, 1:5,000 dil. anti-HDV human serum, or 1:15,000 dil. monoclonal anti-tubulin antibody (Proteintech, Cat# 66031)).

Membranes were washed three times using TBS-T (5 min, RT) and probed for (2 h, RT) with secondary antibody in 1% w/v non-fat milk powder in TBS-T (1:2,000 dil. HRP-coupled goat anti-mouse IgG, 1:2,000 HRP-coupled goat anti-human IgG, or 1:2,000 dil. HRP-coupled goat anti-rabbit IgG in 50% glycerol). Automated detection was performed using SuperSignal West Pico PLUS (Thermo Scientific, #34580) on a ChemiDoc MP imaging system (Bio-Rad).

### Serology

Anti-deltavirus antibodies were detected via immunofluorescence analysis (IFA). HuH7 cells were transfected with DAg-expressing plasmids as described above. The following day, cells were detached using trypsin-EDTA 0.05% and transferred to 8-well chamber slides (Sarstedt, #94.6170.802). At 2 dpt, cells were fixed/permeabilized using ROTI-Histofix with 0.2% Triton-X 100 (15 min, RT). Cells were blocked with PBS containing 10% FBS and then incubated with a 1:20 dil. of animal serum (2 h, 37°C). Cells were washed three times with PBS and then incubated with a 1:500 dil. of goat anti-monkey IgG or goat anti-bat IgG (1 h at 37°C). Following three PBS washes, cells were probed with a 1:100 (bat sera) or 1:400 (monkey sera) dil. of Cy3-coupled donkey anti-goat IgG and 1 µg/ml DAPI (30 min, 37°C). Slides were mounted using Prolong diamond antifade mountant (Invitrogen, #P36970) and cured (24 h, RT, in the dark).

### High-throughput sequencing

High-throughput long-read sequencing was performed to detect potential minority sequence populations containing an edited DAg stop codon. Extracted nucleic acids from original liver tissue were reverse transcribed using Superscript IV (Invitrogen, #18090050) for 20 min at 50°C using 100 nM each of primers HDV-LRS_R1141 and HDV-LRS_R1107 followed by enzyme inactivation for 10 min at 80°C (**Supplementary CTAT Table**). Complementary DNA was subjected to conventional nested PCR using Q5 Hot Start polymerase (New England Biolabs, #M0493S) and the following program: 98°C for 3 min, 35 cycles of (98°C for 10 s, 69°C for 20 s, 72°C for 10 s), 72°C for 2 min. Specific amplification was confirmed by agarose gel electrophoresis. The total PCR reaction was purified using the Monarch Spin PCR and DNA cleanup kit (New England Biolabs, #T1130) and eluted in nuclease-free water. DNA libraries were generated using the Rapid Barcoding kit 96 V14 (Oxford Nanopore Technologies, #SQK-RBK114.96) followed by sequencing on a MinION device. Raw reads were imported into Geneious Prime, trimmed using the BBDuk v38.84 plug-in (minimum read quality of 30 on both ends, discarding trimmed reads of <10 bp length), and resulting reads were mapped against the consensus sequences of RJ-242 and MG-1149, respectively, using the Geneious mapper at highest sensitivity.

## RESULTS

### Detection of deltaviruses

We screened liver specimens of 961 deceased NHP from Brazil for deltaviruses (**table 1**). Two samples (K062_20_RJ-242 and K062_21_MG-1149, hereafter shortened to RJ-242 and MG-1149) tested positive in a broadly-reactive deltavirus-specific nested RT-PCR. Host genera were determined as *Sapajus sp.* (RJ-242) and *Callicebus sp.* (MG-1149) via nested PCR targeting cytochrome C oxidase subunit I as described [19]. The NHP deltaviruses display a highly base-complementary circular structure (**Fig. 1A**), a conserved antisense ORF for DAg, predicted genomic and antigenomic ribozymes, and a high GC content (53.7-54.3%) typical for deltaviruses [20]. Additional putative ORFs of 72-123 amino acids were predicted (**Fig. S2**), of which none were conserved between the two NHP viruses and none were predicted to have transmembrane domains or signal peptides necessary for N-glycosylation, two defining characteristics of viral envelope glycoproteins [21].

**Fig. 1.**
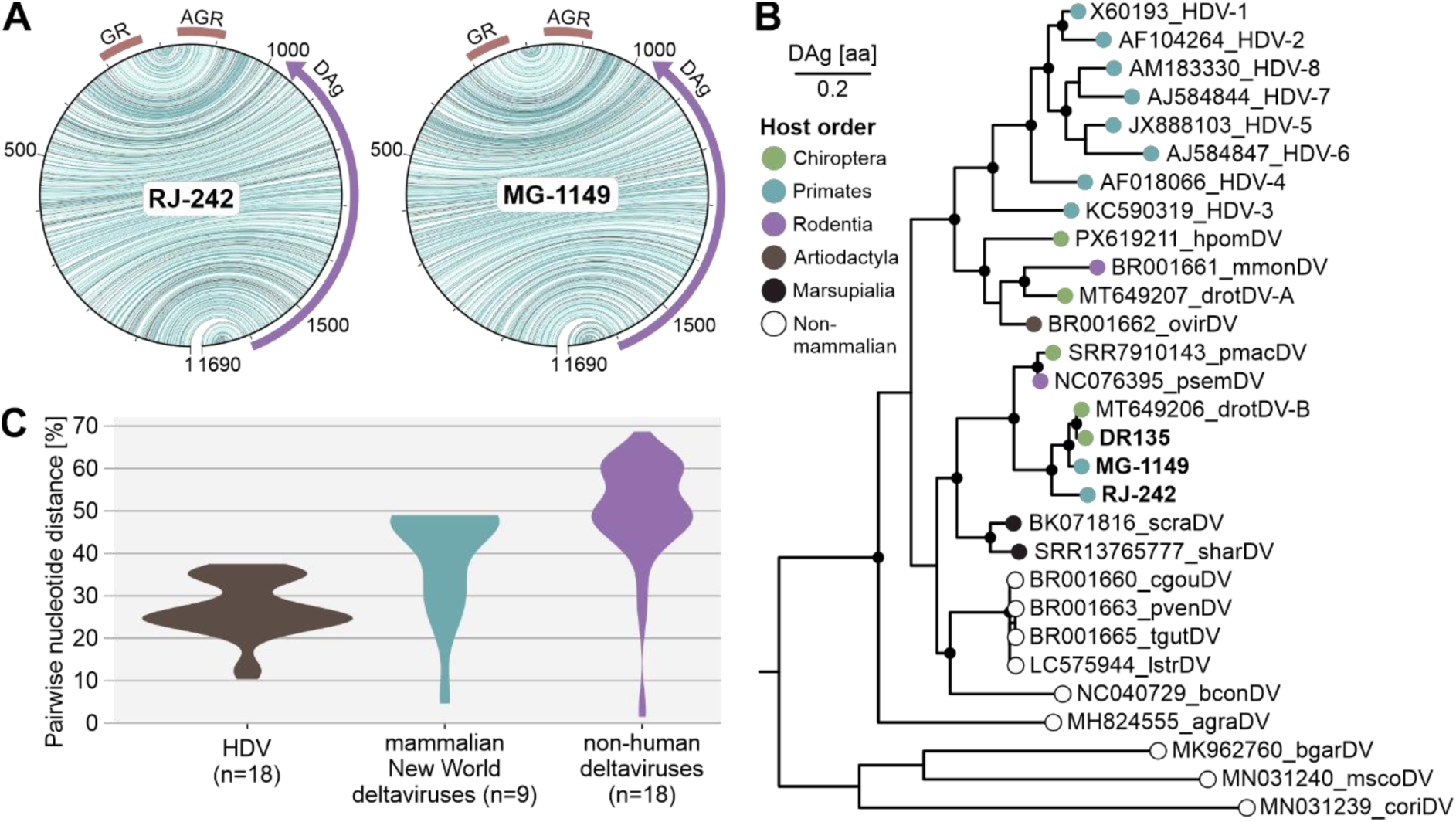
Characterization of non-human primate deltaviruses. **(A)** Base complementary plot based on full genome nucleotide sequences of non-human primate deltaviruses. Base pairings are colored by type of pairing. The open reading frame of delta antigen (DAg; purple), and genomic and antigenomic ribozymes (GR and AGR, respectively; red) are indicated. **(B)** Bayesian phylogeny of vertebrate deltaviruses based on delta antigen amino acid sequence. Tips are colored according to host order. Nodes with posterior probability of >0.8 are indicated with black circles. Termite deltavirus (MK962759) was used as outgroup but removed for graphical clarity. (**C**) Full genome pairwise nucleotide distances of representative sequences of HDV (sub-) species sequences [27], mammalian New World and all non-human vertebrate deltaviruses.

**Table 1:**
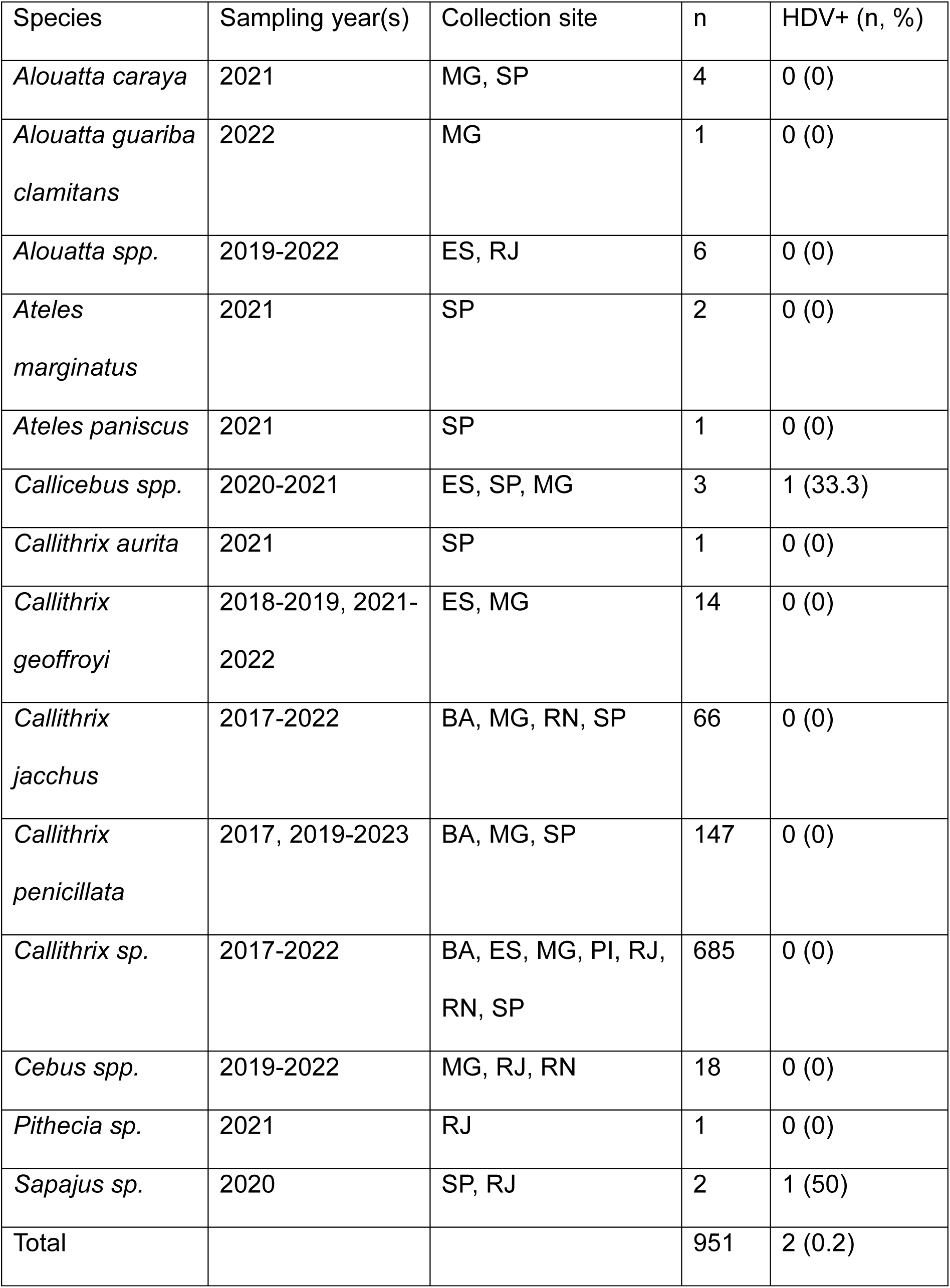
Sample characteristics. Species determination was based on morphology BA: Bahia, MG: Minas Gerais, SP: São Paulo, ES: Espírito Santo, PI: Piauí, RJ: Rio de Janeiro, RN: Rio Grande do Norte.

| Species | Sampling year(s) | Collection site | n | HDV+ (n, %) |
| --- | --- | --- | --- | --- |
| *Alouatta caraya* | 2021 | MG, SP | 4 | 0 (0) |
| *Alouatta guariba clamitans* | 2022 | MG | 1 | 0 (0) |
| *Alouatta* spp. | 2019-2022 | ES, RJ | 6 | 0 (0) |
| *Ateles marginatus* | 2021 | SP | 2 | 0 (0) |
| *Ateles paniscus* | 2021 | SP | 1 | 0 (0) |
| *Callicebus* spp. | 2020-2021 | ES, SP, MG | 3 | 1 (33.3) |
| *Callithrix aurita* | 2021 | SP | 1 | 0 (0) |
| *Callithrix geoffroyi* | 2018-2019, 2021-2022 | ES, MG | 14 | 0 (0) |
| *Callithrix jacchus* | 2017-2022 | BA, MG, RN, SP | 66 | 0 (0) |
| *Callithrix penicillata* | 2017, 2019-2023 | BA, MG, SP | 147 | 0 (0) |
| *Callithrix* sp. | 2017-2022 | BA, ES, MG, PI, RJ, RN, SP | 685 | 0 (0) |
| *Cebus* spp. | 2019-2022 | MG, RJ, RN | 18 | 0 (0) |
| *Pithecia* sp. | 2021 | RJ | 1 | 0 (0) |
| *Sapajus* sp. | 2020 | SP, RJ | 2 | 1 (50) |
| Total |  |  | 951 | 2 (0.2) |

Phylogenetic analysis in a Bayesian framework showed closest relationship of NHP deltaviruses with viruses belonging to Desmodus rotundus deltavirus B (drotDV-B) in a clade that further includes rodent-associated psemDV (**Fig. 1B**). Taxonomically, the new viruses belong to the genus *Thursazvirus*, as DAg amino acid identities were above 60% (75.5-94.4%) (**table S1**) [20].

Full genome pairwise nucleotide distances amongst representative HDV (sub-) species sequences were lower (10.4-37.5%) than between New World mammalian deltaviruses excluding humans (4.6-49.0%) and the entirety of known non-human deltaviruses (1.4-68.7%) (**Fig. 1C**).

### Detection of a hepadnavirus in a saki monkey

To exclude coinfection with a hepadnavirus, we further screened the deltavirus-positive and all other NHP liver specimens from Brazil using a broadly reactive hepadnavirus-specific nested PCR. Screening confirmed the absence of a hepadnavirus in the deltavirus-positive NHP specimens, while a different sample from a saki monkey, *Pithecia sp.* (K062_21_RJ_169, shortened to RJ-169), turned out positive. The complete genome sequence of the saki monkey hepatitis B virus (SMHBV) conformed to hepadnavirus characteristics (**Fig. 2A**) including four ORFs (P, S, X, C). Pairwise sequence comparisons showed high identity to human and NHP hepadnaviruses (79.1-82.5%, **Fig. 2B**). Phylogenetic analysis of full-length hepadnaviral genomes revealed closest relationship with capuchin monkey hepatitis B virus (CMHBV) within the monophyletic primate clade (**Fig. 2C**).

**Fig. 2.**
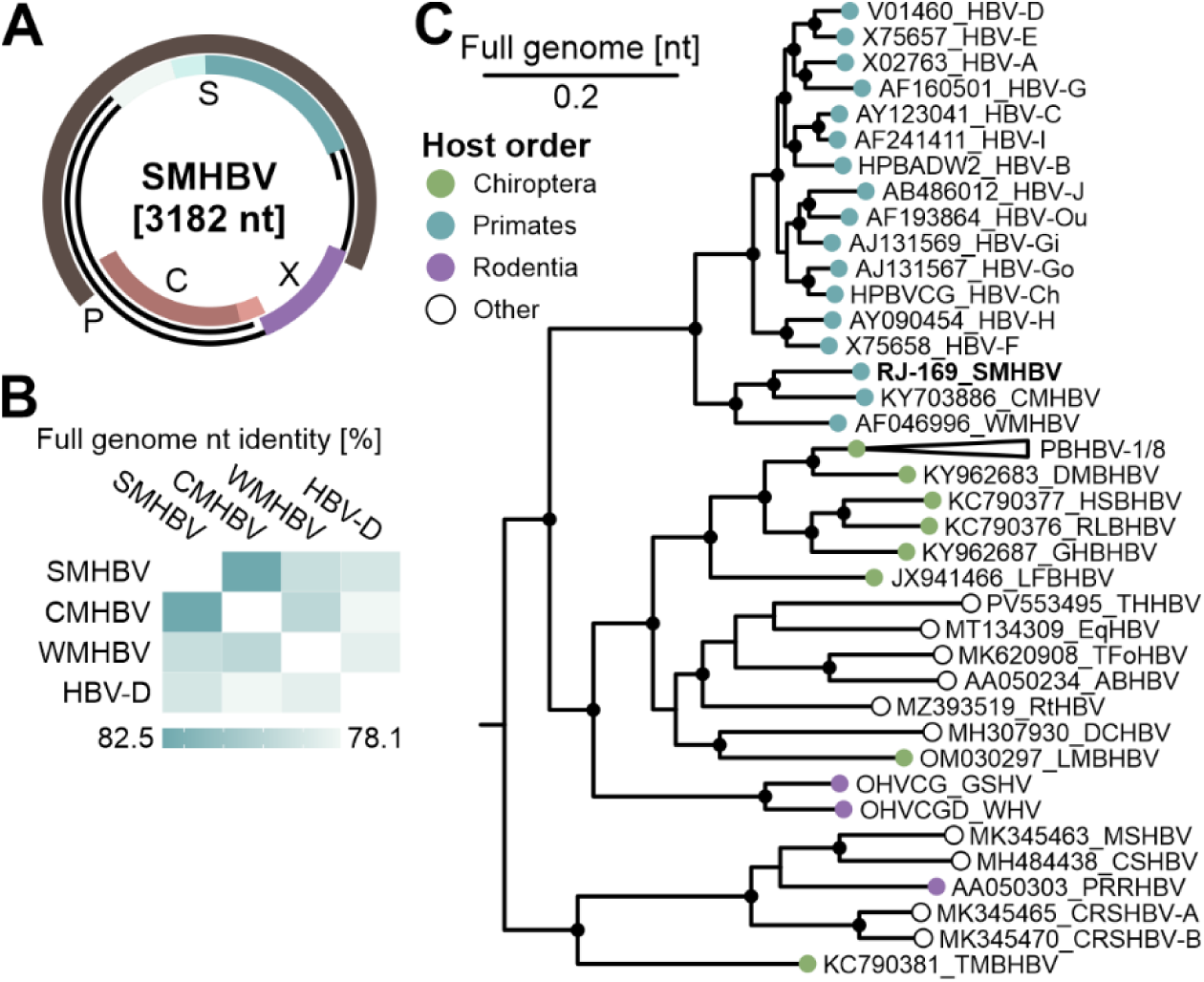
Detection of a saki monkey hepatitis B virus (SMHBV) and putative coinfecting viruses. **(A)** Genome structure of SMHBV. **(B)** Pairwise nucleotide identities of full genome primate hepadnavirus sequences (GenBank acc. no.: V01460 (HBV-D), NC043528 (CMHBV), AF046996 (WMHBV)). **(C)** Bayesian phylogeny of full genome hepadnavirus nucleotide sequences. GenBank or GenBase acc. no. are given in tip names. Tree tips are colored according to host order (Chiroptera (green), Primates (teal), Rodentia (purple), and other (white)). Nodes with posterior probability of >0.95 are indicated with black circles. **(D)** Positive viral families/genera of high-throughput sequencing reads from DNA and RNA sequencing libraries of original liver specimen extracts of RJ-242 and MG-1149. ABHBV: African buffalo hepadnavirus, CMHBV: Capuchin monkey hepadnavirus, CRSHBV-A: Crowned shrew hepadnavirus genotype A, CSHBV: Chinese shrew hepadnavirus, DCHBV: Domestic cat hepadnavirus, DMBHDV: David’s myotis bat hepadnavirus, EqHBV: Equid hepadnavirus, GHBHBV: Greater horseshoe bat hepadnavirus, GSHV: Ground squirrel hepadnavirus, HBV-A: Hepatitis B virus genotype A, HSBHBV: Horseshoe bat hepadnavirus, LFBHBV: long-fingered hepadnavirus, LMBHBV: Large myotis bat hepadnavirus, MSHBV: Musk shrew hepadnavirus, PBHBV-1: Pomona bat hepadnavirus genotype 1, PRRHBV: Pigmy rice rat hepadnavirus, RLBHBV: Roundleaf bat hepadnavirus, RtHBV: Ringtail hepadnavirus, TFoHBV: Tai forest hepadnavirus, THHBV: Tree hyrax hepadnavirus, TMBHBV: Tent-making bat hepadnavirus, WHV: Woodchuck hepadnavirus, WMHBV: Woolly monkey hepadnavirus.

### NHP deltaviruses replicate without the expression of a large delta antigen

HuH7 cells transfected with genomic dimer plasmids of RJ-242 and MG-1149 showed increasing expression of intracellular DAg over time via immunoblot (**Fig. 3A-B**). The dimer plasmids containing the R13A mutation in DAg, inhibiting antigenome to genome conversion [18], showed a decreasing trend of DAg expression linked to the fading transcription from transfected plasmid DNA (**Fig. 3A-B**). A second, larger protein corresponding to L-DAg was detected in wildtype HDV-transfected HuH7 cells from 6 dpt (**Fig. 3C**), but neither in HDV-R13A-(**Fig. 3C**), nor NHP deltavirus-transfected cells (**Fig. 3A-B**), despite encoding putative C-terminal extensions (**Fig. S1B**). A time-dependent decrease in DAg signal was also observed after transfection of a cytomegalovirus promoter-controlled DAg-expressing plasmid (**Fig. 3D**).

**Fig. 3.**
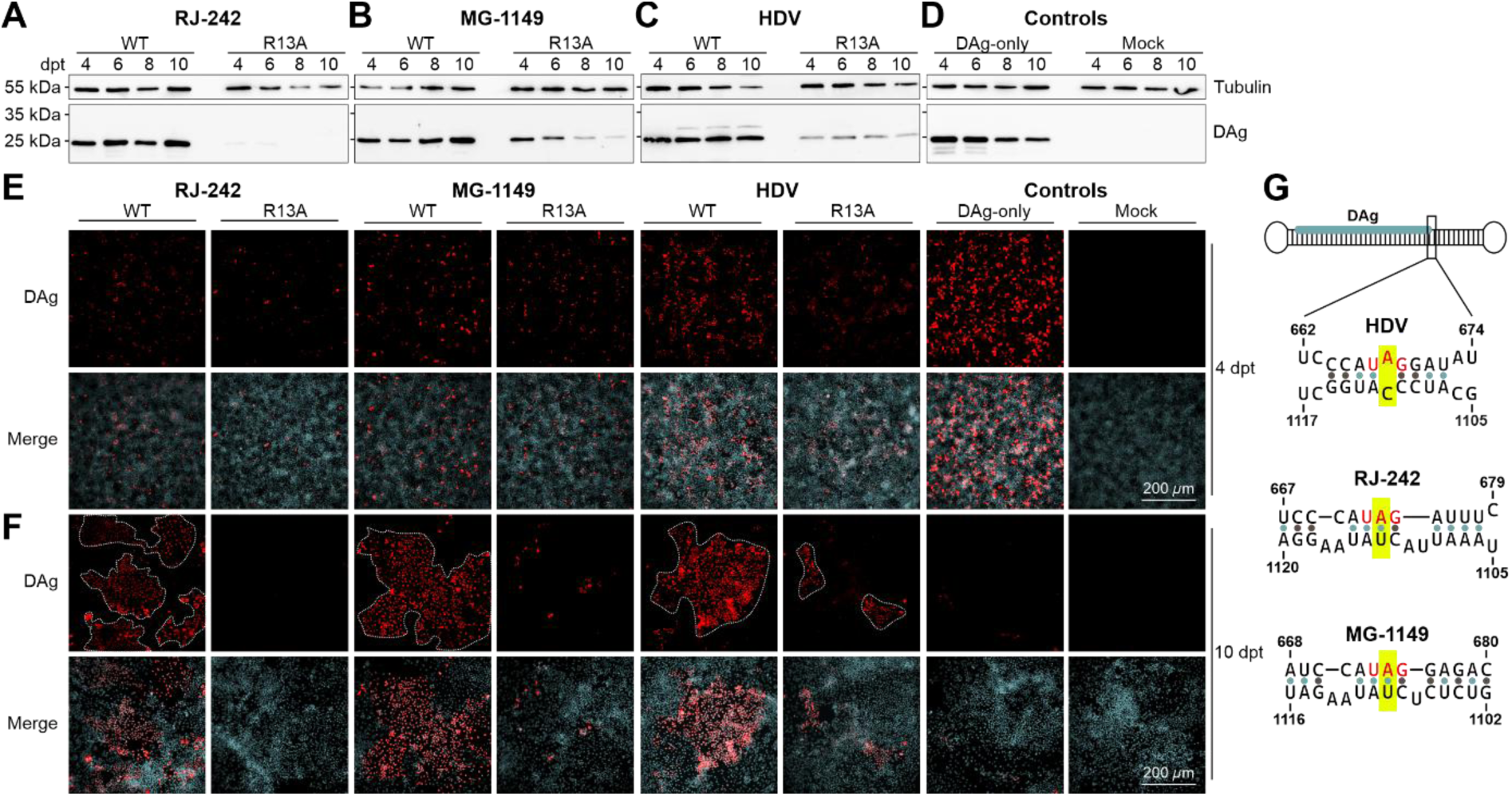
Non-human primate deltaviruses replicate without large delta antigen (DAg) *in vitro*. DAg immunoblots of HuH7 cells transfected with the RJ-242 dimer **(A**), MG-1149 dimer **(B)**, hepatitis D virus (HDV) dimers **(C)**, or control plasmids **(D)** lysed at the indicated timepoints (4 to 10 days post transfection (dpt)). Cell division-mediated spread experiments of transfected HuH7 cells **(E)** 4 dpt unsplit and **(F)** 10 dpt following a 1:100 split on 1 dpt. **(G)** *In silico* RNA folding of deltaviral antigenomes. The amber stop codon triplet of DAg is shown in red. Numbering according to the respective genome positions. The ICTV reference sequence of HDV-1 was used (GenBank accession number: AF104263).

Amplicon-based long-read sequencing of original liver tissue extracts was also not indicative of an editing event affecting the DAg stop codon. There were no reads after quality filtering containing an adenosine-guanosine-transition in that position (RJ-242 (A: 872, T: 1, G: 0, C: 0) and MG-1149 (A: 750, T: 0, G: 0, C: 0)), in contrast to a 60:40 split between unedited and edited genomes described for HDV [22].

In line with the sequencing data, the unpaired adenosine at position 668 within the DAg amber stop codon of the human HDV antigenome (numbering according to reference genome, GenBank accession number: AF104263) is essential for efficient ADAR-directed editing that enables L-DAg expression [23]. *In silico* RNA folding of the complete deltaviral antigenomes revealed the analogous adenosine to be base paired in both NHP deltaviruses (**Fig. 3G**). Taken together, the molecular and computational data exclude the expression of an L-DAg in the novel NHP deltaviruses.

To investigate whether the NHP deltaviruses can be transmitted to descendent cells during mitosis described for other deltaviruses [17,24], cell division-mediated spread experiments were performed. HuH7 cells transfected with wildtype and R13A-mutant dimer plasmids were highly diluted on 1 dpt and grown until 10 dpt. While there are individual DAg-positive cells in unsplit wells (grown until 4 dpt) in all conditions except mock control (**Fig. 3E**), large DAg-positive cell clusters appear only after transfection of wildtype dimers on 10 dpt following high dilution (**Fig. 3F**; white dashed lines).

### Geographically widespread vampire bat populations carry highly related deltaviruses

The presence of viruses in common vampire bats closely related to the novel NHP deltaviruses prompted us to screen vampire bat specimens previously collected [10]. We screened a total of 448 liver and 229 spleen specimens from Brazil and Costa Rica, respectively. While all Brazilian specimens tested negative in RT-PCR, we were able to recover a complete genome from a Costa Rican vampire bat (K083_DR135, shortened to DR135). This vampire bat deltavirus was closely related to previously described viruses found in Peruvian vampire bats [7] belonging to drotDV-B (**Fig. 4A**), despite a geographic distance of ∼2800 km. Realtime RT-PCR of available organ specimens detected deltavirus RNA only in liver and spleen (2.9 × 10^4^ and 6.8 × 10^3^ copies/g, respectively) (**Fig. 4B**).

**Fig. 4.**
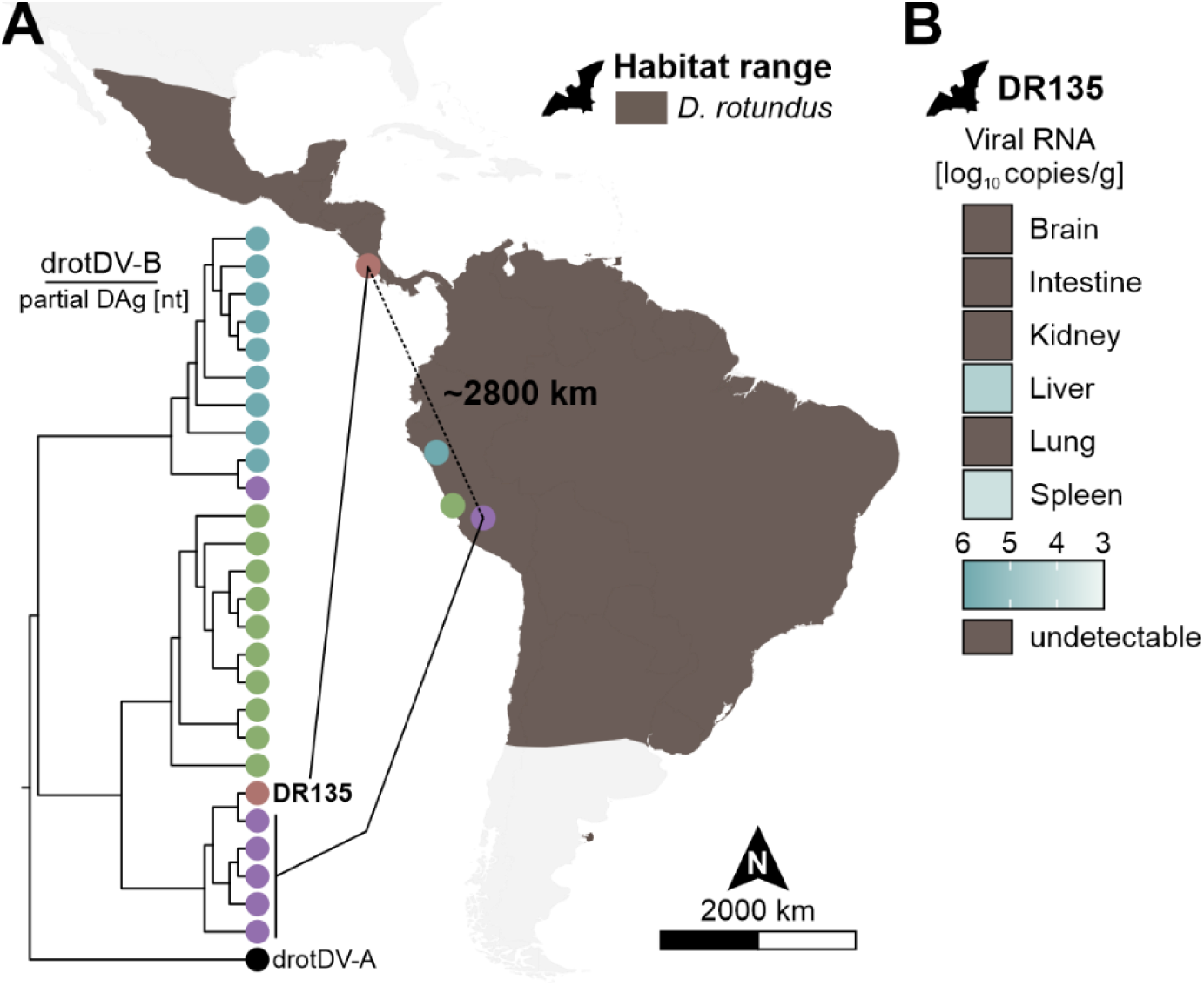
Characterization of vampire bat deltavirus DR135. **(A)** Cladogram of a Bayesian phylogeny of partial delta antigen nucleotide sequences (213 nucleotides) of DR135 and sequences of Desmodus rotundus deltavirus B (drotDV-B) published in Bergner *et al.* [7]. DrotDV-A (GenBank accession number: MT649207) was used as outgroup. Corresponding sampling sites and tree tips are colored. Habitat range of the common vampire bat (*Desmodus rotundus*) is based on IUCN Red List data. **(B)** Deltaviral RNA load in organ specimens of common vampire bat DR135 by real-time RT-PCR.

### High prevalence of deltavirus-specific antibodies in sera of bats but not non-human primates

First, we performed IFA based on NHP DAg-expressing HuH7 cells. We tested a total of 249 NHP sera from Brazil (160 *Callithrix sp.*, 84 *Sapajus sp.*, 4 *Lagothrix lagotricha*, 1 *Callicebus sp.*), of which none showed reactivity with NHP DAg. A human anti-HDV serum could be detected using the identical setup of secondary and tertiary antibodies indicating robust assay conditions (**Fig. 5A**). We further performed serological testing of 112 *Desmodus rotundus* sera from Costa Rica with DR135 DAg-expressing HuH7 cells and detected deltavirus-specific antibodies in seven sera (6.2%, 95% CI: 1.8-10.7) (**Fig. 5B**) with endpoint titers between 1:40 and 1:5,120 (**Table S2**). Notably, the serum of the deltavirus RNA-positive bat (DR135) did not show DAg-reactivity in IFA.

**Fig. 5.**
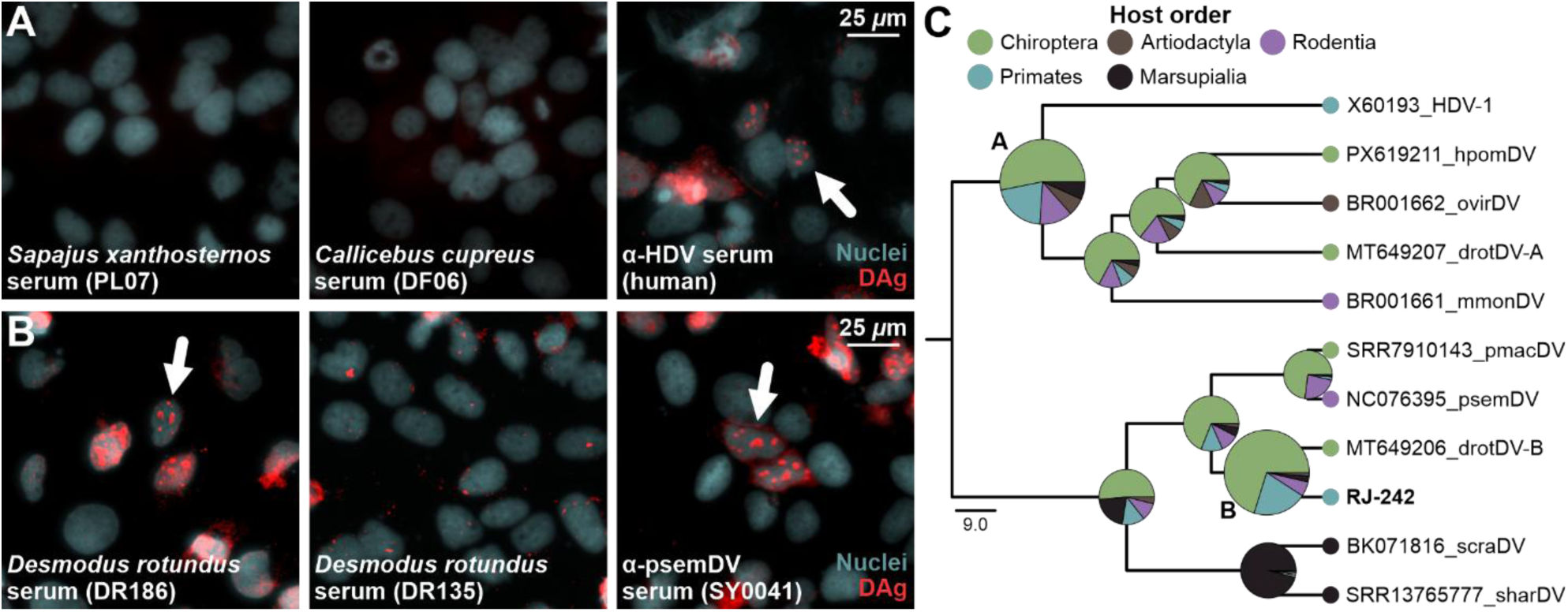
Putative origin of non-human primate (NHP) deltaviruses in bats. **(A)** Exemplary NHP deltavirus-derived delta antigen (DAg)-based immunofluorescence analyses of two non-reactive NHP sera and a cross-reactive anti-HDV antiserum. Characteristic nuclear DAg staining (red) [1] is marked with white arrows. **(B)** Exemplary DR135-derived DAg-based immunofluorescence analyses of common vampire bat sera. A cross-reactive anti-Proechimys semispinosus deltavirus (psemDV) rabbit antiserum (SY0041) was used as positive control. **(C)** Ancestral state reconstruction of mammalian deltaviruses by host order performed with BEAST. Node ancestral likelihoods in percent are colored by host order. The GenBank/SRA accession numbers are stated.

### A putative origin of primate deltaviruses in bats

The high prevalence of anti-deltavirus antibodies in bats but not NHP despite phylogenetic relatedness of their viruses hinted at the former to represent a deltavirus reservoir for primates. We performed Bayesian ancestral state reconstruction (ASR) of mammalian deltaviruses to determine the likelihood of primate deltaviruses to have originated in bats. The analysis favored a bat over a primate origin for the two clades containing human and NHP deltaviruses, respectively (**Fig. 5C**, node A (53.2% bat vs. 20.1% primate origin) and node B (70.3% bat vs. 20.6% primate origin)).

## CONCLUSIONS

The two novel NHP deltaviruses share common features of deltaviruses including high self-complementarity and expression of a DAg. *In vitro* data were compatible with cell division-mediated persistence [20] independent of a large isoform of DAg, comparable to other non-human deltaviruses [1,17].

The NHP deltaviruses were closely related to deltaviruses from vampire bats and belonged to a clade different from HDV. The absence of anti-deltavirus antibodies in NHP opposed sustained circulation of these deltaviruses in the NHP population in Brazil, when assuming a seroconversion rate similar to humans, where anti-HDV antibodies develop in all immunocompetent individuals infected [25]. In consequence, NHP deltaviruses described here may derive from a bat-associated source, likely from bats, as antibody detection rate in tested vampire bat sera was high (6.2%, 1.8-10.7). The antibody detection rate in this study was lower than for the related psemDV in rodents, where 13.7% of animals overall were antibody-positive (and 57% of RNA-positive animals) [17], likely attributable to the ecological confinement of the investigated rodent population. This bat-to-primate spillover scenario is consistent with the putative bat origin of NHP deltaviruses (and HDV) by ASR, highlighting the necessity to recover more deltaviral sequences from bats and NHP to resolve the evolutionary history of these two clades. While deltavirus spillovers should be considered rare, introduction of a deltavirus ancestral to HDV from bats into humans is conceivable. Particularly vampire bats may represent a significant deltavirus reservoir for primates as high biting rates of humans (>20%) have been reported [26] and drotDV-B, the closest known relative of the NHP deltaviruses, has been detected in vampire bat saliva [7]. However, the data on primate deltavirus origins must beinterpreted with caution as the current number and diversity of deltaviruses is likely limited by significant undersampling and future discoveries may adversely affect macroevolutionary reconstructions.

In conclusion, the missing link remains a non-human deltavirus genetically precursory to (or already capable of) L-DAg expression. Hence, continued large-scale screening of New World NHP and reservoir hosts (i.e., bats) using targeted screening assays is crucial to solve the riddle of HDV emergence. This study’s data are consistent with deltaviral cross-order host shifts, highlighting the susceptibility of primates to reservoir-bound deltaviruses and providing a biologically plausible scenario for the emergence of HDV in humans.

## ACKNOWLEDGEMENTS

We thank Sebastian Brünink, Patricia Tscheak and Anja Kühl (from iPATH.Berlin at Charité - Universitätsmedizin Berlin) for technical assistance. We thank Eduardo Martins Netto, Carlos Brites, Celia Pedroso, Andreas Stöcker, Cecília Kierulff, Márcio Port-Carvalho, Tiago Ferreira da Silva, Ivone Kuribara, Hélio Ramos, and the teams at the Wildlife Triage and Rehabilitation Center (CETRAS-SP, Tietê Ecological Park), the Sagui Legal Commercial Breeding Facility, Instituto Florestal, and the Military Police of the State of São Paulo for their assistance and collaboration during fieldwork and sample collection.

## Author contributions

FL: conceptualization, formal analysis, investigation, validation, data curation, writing - original draft, visualization; AMS: conceptualization, formal analysis, investigation, validation, data curation, visualization; MAMG: resources, investigation; AAA: investigation, visualization; CVM: investigation, resources; IOV: investigation, resources; BFdCDS: investigation, resources; AJBC: resources; GHM: resources; DG: resources; ACdAC: resources, supervision; CRF: resources, supervision; ELD: resources, supervision; AMBdF: resources, supervision; JFD: conceptualization, resources, writing – review and editing, supervision, funding acquisition.

## Data availability statement

Deltavirus consensus sequences described in this study are available under GenBank accession numbers XYZ-XYZ. Cytochrome B (CYTB) and cytochrome C oxidase subunit I (COI) consensus sequences of deltavirus-positive animals are available under GenBank accession numbers XYZ-XYZ. Filtered reads of untargeted short-read and amplicon-based long-read sequencing were deposited in SRA (Bioprojects XYZ and XYZ, respectively). All other original data are available from the corresponding author upon reasonable request.

