## Supplementary data for "Infection of neotropical non-human primates with bat deltaviruses"

This file contains:

Figures S1-S2

Tables S1-S2

Supplementary references

### Supplementary figures

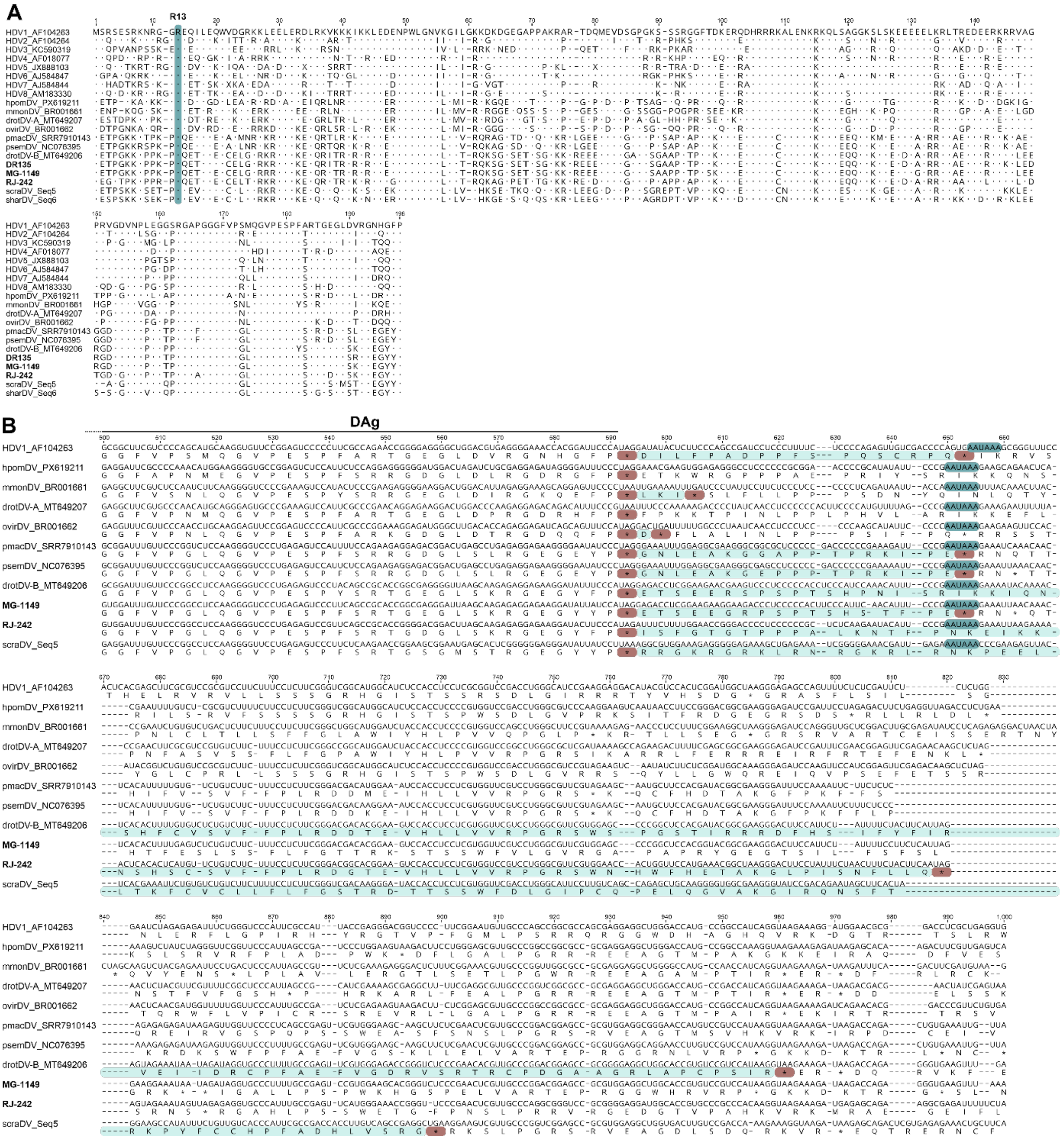

**Fig. S1. Delta antigen (DAg) alignments and putative large DAg extensions.** (A) Amino acid alignment of DAg of mammalian kolmioviruses. The residue R13 (turquoise) is methylated and essential for antigenome to genome conversion [1]. (B) Nucleotide alignment of mammalian kolmioviruses with translation of the DAg open reading frame. Stop codons of the small and putative large DAg (red), putative large DAg extension (light blue), poly-adenylation signal (turquoise) are indicated.

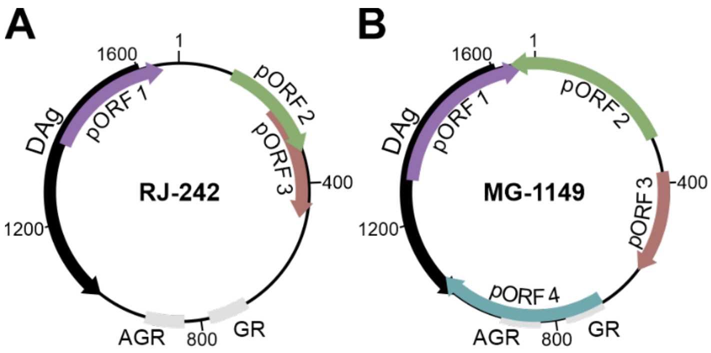

**Fig. S2. Putative open reading frames (pORF) in the non-human primate kolmioviruses.** Open reading frames of RJ-242 (A) and MG-1149 (B) were predicted using Geneious Prime's "Find ORFs" feature with a minimum length of 200 nucleotides and an ATG start codon only. AGR, antigenomic ribozyme; DAg, delta antigen; GR, genomic ribozyme.

**Supplementary tables**

**Table S1. Pairwise amino acid identities [%] of delta antigens of mammalian hosts.** GenBank/SRA accession numbers: drotDV-A (MT649207), drotDV-B (MT649206), HDV (AF104263), hpomDV (PX619211), ovirDV (BR001662), mmonDV (BR001661), pmacDV (SRR7910143), psemDV (NC076395), scraDV (BK071816), sharDV (SRR13765777).

| | **1** | **2** | 3 | 4 | 5 | 6 | 7 | 8 | 9 | 10 | 11 | 12 |
|---|---|---|---|---|---|---|---|---|---|---|---|---|
| **1. RJ-242** | | 84.7 | 84.7 | 75.5 | 77.0 | 67.2 | 68.7 | 57.2 | 52.3 | 58.8 | 53.9 | 52.3 |
| **2. MG-1149** | 84.7 | | 94.4 | 78.6 | 80.6 | 69.7 | 70.8 | 57.7 | 52.3 | 59.3 | 58.0 | 52.8 |
| 3. drotDV-B | 84.7 | 94.4 | | 78.1 | 80.1 | 68.2 | 70.3 | 58.2 | 51.3 | 58.2 | 57.5 | 51.3 |
| 4. pmacDV | 75.5 | 78.6 | 78.1 | | 95.9 | 69.2 | 67.7 | 56.7 | 54.4 | 56.2 | 56.0 | 49.7 |
| 5. psemDV | 77.0 | 80.6 | 80.1 | 95.9 | | 70.8 | 70.3 | 57.7 | 54.9 | 55.2 | 55.4 | 50.8 |
| 6. scraDV | 67.2 | 69.7 | 68.2 | 69.2 | 70.8 | | 86.2 | 60.1 | 56.3 | 56.5 | 55.7 | 51.5 |
| 7. sharDV | 68.7 | 70.8 | 70.3 | 67.7 | 70.3 | 86.2 | | 58.0 | 55.7 | 58.0 | 57.3 | 51.5 |
| 8. ovirDV | 57.2 | 57.7 | 58.2 | 56.7 | 57.7 | 60.1 | 58.0 | | 73.6 | 74.7 | 67.4 | 59.3 |
| 9. hpomDV | 52.3 | 52.3 | 51.3 | 54.4 | 54.9 | 56.3 | 55.7 | 73.6 | | 68.4 | 62.7 | 58.0 |
| 10. drotDV-A | 58.8 | 59.3 | 58.2 | 56.2 | 55.2 | 56.5 | 58.0 | 74.7 | 68.4 | | 70.5 | 62.4 |
| 11. mmonDV | 53.9 | 58.0 | 57.5 | 56.0 | 55.4 | 55.7 | 57.3 | 67.4 | 62.7 | 70.5 | | 56.5 |
| 12. HDV | 52.3 | 52.8 | 51.3 | 49.7 | 50.8 | 51.5 | 51.5 | 59.3 | 58.0 | 62.4 | 56.5 | |

Abbreviations: drotDV, Desmodus rotundus [common vampire bat] deltavirus; HDV, hepatitis D virus; hpomDV, Hipposideros pomona [Pomona roundleaf bat] deltavirus; ovirDV, Odocoileus virginianus [white-tailed deer] deltavirus; mmonDV, Marmota monax [woodchuck] deltavirus; pmacDV, Peropteryx macrotis [lesser dog-like bat] deltavirus; psemDV, Proechimys semispinosus [Tome's spiny rat] deltavirus; scraDV, Sminthopsis crassicaudata [fat-tailed dunnart] deltavirus; sharDV, Sarcophilus harrisii [Tasmanian devil] deltavirus.

**Table S2. Anti-deltavirus immunofluorescence analysis endpoint titers of reactive *Desmodus rotundus* sera.** Immunofluorescence analysis based on K083_DR135-derived delta antigen (DAg)-expressing HuH7 cells.

| Sample | DR042 | DR044 | DR052 | DR098 | DR132 | DR168 | DR186 |
|---|---|---|---|---|---|---|---|
| IFA endpoint titer | 1:5,120 | 1:2,560 | 1:80 | 1:5,120 | 1:5,120 | 1:40 | 1:1,280 |
